# Development of PAR-CLIP to analyze RNA-protein interactions in prokaryotes

**DOI:** 10.1101/855189

**Authors:** Sandeep Ojha, Chaitanya Jain

## Abstract

The ability to identify RNAs that are recognized by RNA-binding proteins (RNA-BPs) using techniques such as “Crosslinking and Immunoprecipitation” (CLIP) has revolutionized the genome-wide discovery of RNA targets. Among the different versions of CLIP developed, the incorporation of photoactivable nucleoside analogs into cellular RNA has proven to be especially valuable, allowing for high efficiency photoactivable ribonucleoside-enhanced CLIP (PAR-CLIP). Although PAR-CLIP has become an established technique for use in eukaryotes, it has not yet been applied in prokaryotes. To determine if PAR-CLIP can be used in prokaryotes, we first investigated whether 4-thiouridine (4SU), a photoactivable nucleoside, can be incorporated into *E. coli* RNA. After determining 4SU incorporation into RNA, we developed suitable conditions for crosslinking of proteins in *E. coli* cells and for the isolation of crosslinked RNA. Applying this technique to Hfq, a well-characterized regulator of small RNA (sRNA) - messenger RNA (mRNA) interactions, we showed that PAR-CLIP identified most of the known sRNA targets of Hfq. Based on our results, PAR-CLIP represents an improved method to identify the RNAs recognized by RNA-BPs in prokaryotes.

## Introduction

RNA-BPs constitute a large and diverse class of proteins that regulate nearly every aspect of RNA function inside the cell. Among the hundreds of such proteins, the specific functions of many are not well understood. An important component of defining the cellular roles of RNA-BPs is to identify the RNAs they interact with inside the cell. Such efforts are not only useful to define RNA-BP function, but also the basis of RNA recognition through a systematic analysis of their RNA targets.

The first genome-wide method developed to identify the specific RNAs recognized by RNA-BPs was based on RNA immunoprecipitation (RIP) ^1^. In this method, the protein of interest, as well as its associated RNAs, are purified by using anti-RNA-BP antibodies. Thereafter, the RNAs are extracted and their identities are revealed either by sequencing or by hybridization to DNA microarrays. Over the past few years RIP has become largely supplanted by CLIP, a method that is conceptually similar to RIP, except that the RNA-BPs are first crosslinked to their RNA targets *in vivo* ^2,3^. The development of CLIP stemmed, in part, from concerns that the non-covalent nature of protein-RNA interactions in RIP could result in the loss of many bound RNAs, especially if extensive purification steps are employed.

Two different procedures for crosslinking have been applied for CLIP. In the first, a chemical crosslinker, such as formaldehyde, has been used ^4,5^. In the second approach, RNA-protein crosslinks are generated by irradiating cells with short-wavelength ultraviolet (UV) light ^2,3^. Despite widespread adoption, each method suffers from limitations that can reduce their effectiveness. With chemical crosslinking, a significant drawback is that the crosslinking reagents induce not only protein-RNA, but also protein-DNA and protein-protein crosslinks. Therefore, the RNAs captured can include those that are bound by other RNA-binding proteins that interact with the RNA-BP of interest. In contrast, the efficiency of crosslinking is low with standard UV-CLIP, which can impact the coverage of RNA targets identified using this method. Any improvements to these techniques, therefore, are expected to yield a more complete and accurate picture of RNA-BP targets inside the cell.

To improve on existing procedures, a new method, PAR-CLIP, was developed ^6^. In this method, cells are grown in the presence of a photoactivable nucleotide added to the growth medium, the most commonly used being 4-thiouridine (4SU). Incorporation of 4SU into RNA, followed by irradiation with long-wavelength UV light, has been reported to enhance crosslinking efficiency by 100-1000-fold relative to conventional CLIP ^6^. An additional advantage of PAR-CLIP is that crosslinked 4SU residues are frequently misread as cytosine residues during the reverse transcription step of sequence library preparation ^6–8^. Consequently, an analysis of T to C conversion frequency in the sequence data facilitates the identification of the specific sites of RNA-protein interactions, accelerating the process of identifying the RNA recognition motifs of RNA-BPs.

Although PAR-CLIP is now an established procedure in eukaryotes, to the best of our knowledge, it has not yet been implemented in prokaryotes. To determine the feasibility of applying this technique to prokaryotic organisms, we first investigated whether 4SU can be incorporated into RNA in *E. coli.* After demonstrating 4SU incorporation, we then applied PAR-CLIP to Hfq, a well-characterized regulator of sRNA-mRNA interactions, and found that PAR-CLIP effectively identifies most of the sRNAs previously shown to be associated with Hfq. Based on our results, we anticipate that the use of PAR-CLIP to prokaryotes will provide a superior method to identify the *in vivo* RNA targets of RNA-BPs in such organisms.

## Results

### 4SU is incorporated into RNA in E. coli

To determine whether 4SU can be incorporated into *E. coli* RNA, a wild-type strain (CJ2109) was grown in LB medium or LB supplemented with 300 μM 4SU. RNA was isolated from mid-log cultures and digested with nucleases and phosphatases to nucleosides. The digested RNA was analyzed by high performance liquid chromatography (HPLC) with readings at 260 nm (all nucleotides) or 330 nm (4SU). Both sets of RNAs gave similar peaks for the standard nucleosides (Fig. 1). However, with detection at 330 nm, the RNA samples isolated from cells grown with 4SU yielded a significantly higher signal compared to cells grown without 4SU. Based on peak heights and a comparison with defined nucleoside standards, we calculated that 1.3% of uridine residues in the latter RNA sample were replaced with 4SU. These experiments indicated that 4SU can be incorporated into RNA *in vivo*, further implying that *E. coli* RNA polymerase does not discriminate against the incorporation of 4SU into RNA during transcription.

**Figure. 1.**
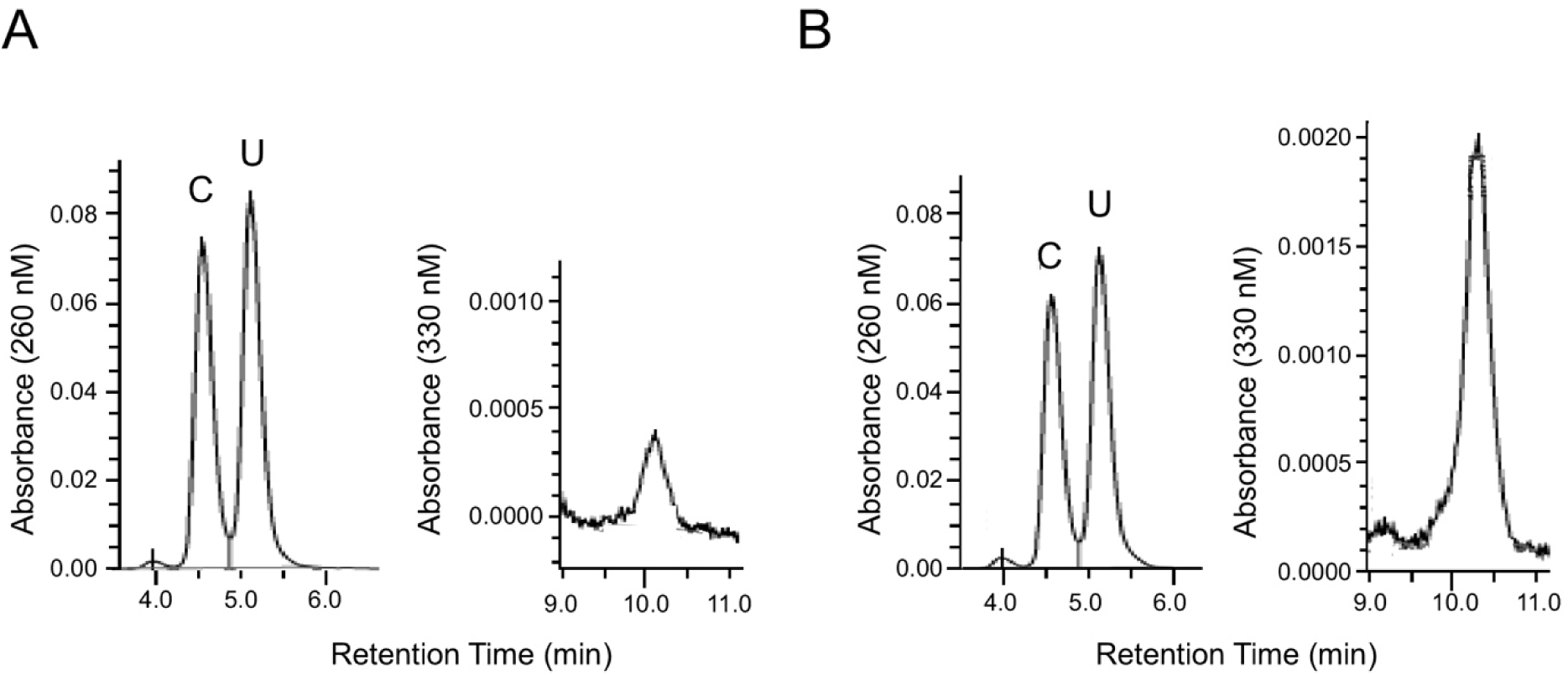
HPLC analysis. Total cellular RNA was isolated from cells grown in the absence (A) or presence of 300 μM 4SU (B), enzymatically digested to nucleosides, and the products were resolved by HPLC with detection at 260 nm or 330 nm. Only the regions corresponding to the retention times for uracil (U) and 4SU are shown. For the 260 nm trace, the mutually overlapping cytosine (C) peak is also included. A background 330 nm absorbance of unknown origin is present in the RNA sample isolated from cells grown without 4SU.

### PAR-CLIP of FLAG-tagged proteins

To determine whether 4SU containing RNA can be crosslinked to RNA-BPs, CJ2109 was transformed with pCJ1099, a plasmid that encodes the DEAD-box protein RhlE tagged with a 3X FLAG epitope at its C-terminal end. We have previously shown that RhlE binds to the 50S ribosomal subunit and to 70S ribosomes ^9^. The transformed strain (CJ2191) was grown to midlog phase in the absence or the presence of different concentrations of 4SU in the medium, followed by harvesting of cells and irradiation with long-wavelength UV light (365 nm). The cells were lysed, incubated with beads coupled with anti-FLAG antibodies, and unbound proteins were removed through multiple washes. Thereafter, the crosslinked RNAs were fragmented on beads and labeled at the 5’ ends with *γ*-^32^P-ATP. The labeled proteins were eluted from beads, fractionated by using polyacrylamide gel electrophoresis (PAGE), transferred to nitrocellulose, and exposed to a phosphorimager screen to visualize the crosslinked RNAs. Following the exposures, the blots were re-used for western-blot analysis using anti-FLAG antibodies.

A comparison of the signal derived from cells grown in the presence of different amounts of 4SU indicated that low levels of radiolabeled RNA were crosslinked to RhlE when cells were grown in the absence or the presence of 100 μM 4SU (Fig. 2A). Significant amounts of label were observed for cells grown with 250 μM 4SU and a further signal increase was observed with 500 μM 4SU. These results show that the efficacy of RNA crosslinking is dependent upon the concentration of 4SU in the medium. Surprisingly, when cells were grown with 1 mM 4SU, the signal was found to be reduced. However, we noticed those cells exhibited slower growth, suggesting that the reduced efficiency of 4SU incorporation into RNA could be related to growth inhibition.

**Figure. 2.**
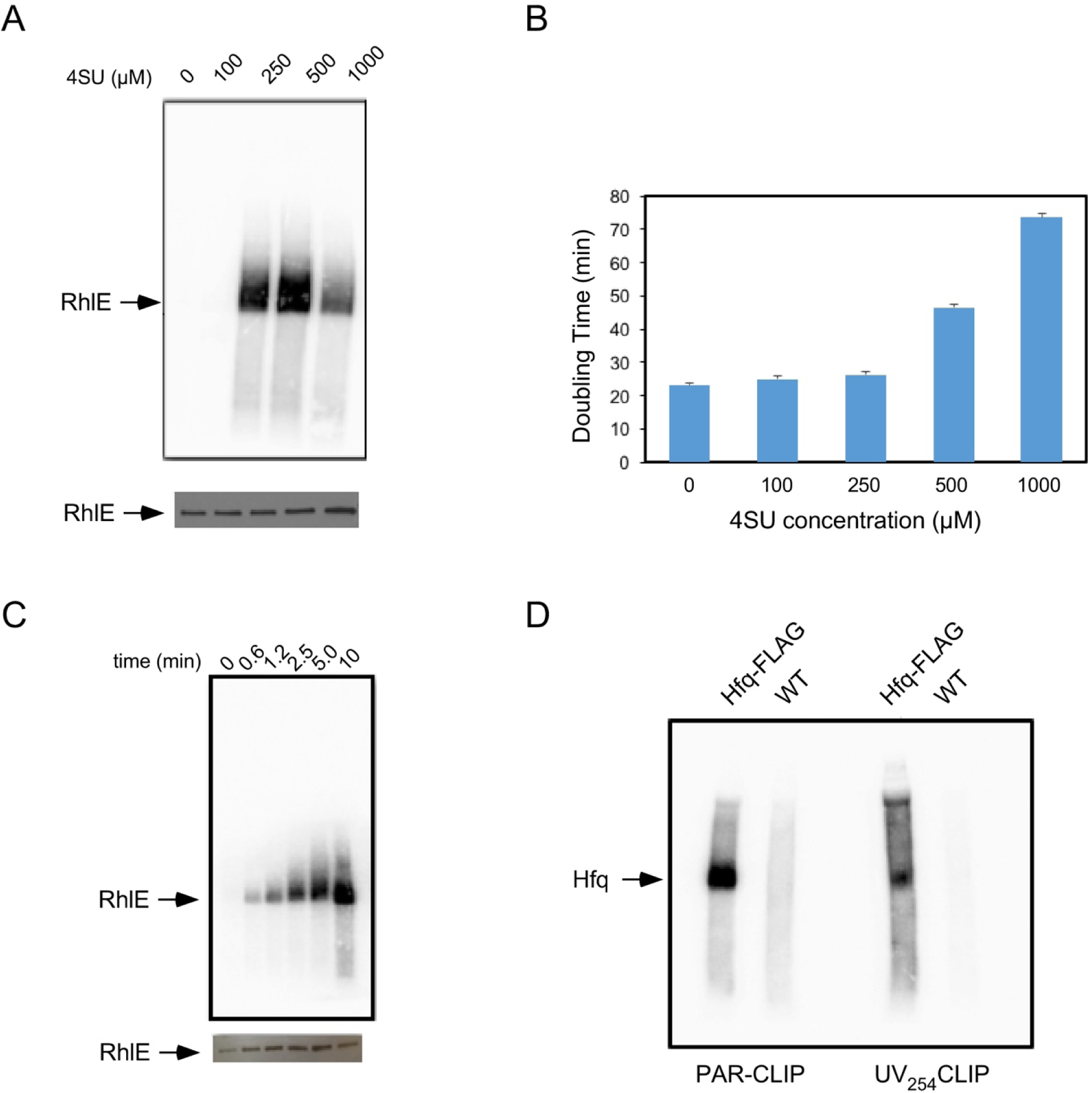
PAR-CLIP on FLAG-tagged proteins. (A) CJ2191 was grown in LB supplemented with different concentrations of 4SU and cells were crosslinked at 365 nm after harvesting. FLAG-tagged RhlE was purified by using anti-FLAG antibodies and the RNAs crosslinked to RhlE were radiolabeled. Proteins were resolved by gel electrophoresis and transferred to nitrocellulose filter. Top, the nitrocellulose filter was exposed to a phosphorimager screen to evaluate the extent of RNA crosslinking to RhlE; bottom, the filter was subsequently analyzed by Western blot using anti-FLAG antibodies. (B) Effect of 4SU on cell growth. CJ2109 was grown in LB supplemented with the indicated concentrations of 4SU and cell doubling times were determined. (C) The effect of UV irradiation time on cross-linking efficiency. CJ2191 cells were irradiated at 365 nm for different periods of time, and crosslinked RhlE was analyzed as in (A). Top, phosphorimager scan; bottom, western-blot analysis. (D) Comparison of PAR-CLIP with UV-CLIP. Strains CJ2192 (Hfq-FLAG) or CJ2109 (WT) were grown in the presence or absence of 250 μM 4SU and irradiated after harvest with 365 nm or 254 nm UV light for five minutes, respectively. FLAG-tag protein purification and RNA labeling was performed as in (A).

To determine the relationship between 4SU concentration and cell growth, cells were grown in LB supplemented with different concentrations of 4SU and cell growth was monitored (Fig. 2B). Little or no growth inhibition was observed with 250 μM 4SU or less, but higher 4SU concentrations were found to slow growth and increase the cell doubling time. These results suggest that the reduced crosslinking observed at 1 mM 4SU concentration (Fig. 2A) might be caused by effects on cell physiology, such as an inhibition of 4SU incorporation into RNA. In order to minimize such effects, subsequent studies were performed on cells grown with 250 μM 4SU.

To further optimize the crosslinking efficiency of 4SU-containing RNAs, cells were irradiated with 365 nm UV light for different amounts of time (Fig. 2C). Increasing the irradiation time for up to 10 minutes resulted in increased levels of crosslinking, indicating that the extent of crosslinking was not saturated even with the longest irradiation time used. Finally, we compared the efficacy of PAR-CLIP with standard UV CLIP. With an intention to generate data for a well characterized RNA-BP, we constructed a strain (CJ2192) that expresses the Hfq protein tagged with a FLAG-epitope at its C-terminus. Hfq is a well-studied regulator of gene expression, and multiple studies have shown that it binds to several regulatory sRNAs inside the cell ^10,11^. CJ2192 was grown either in the presence or the absence of 4SU, and the cells were crosslinked at 365 nm or 254 nm, respectively. To determine whether there is any non-specific background, the same procedure was repeated on the untagged strain (CJ2109). After crosslinking, the crosslinked RNAs were analyzed as described above. With both PAR-CLIP and standard CLIP, a signal corresponding to Hfq could be observed (Fig. 2D). In addition, some background both below and above the main band was clearly observed. The origin of this background is unclear, but it could be due to free RNA or to other RNA-BPs that bind to the antibody-coupled beads ^12^. With PAR-CLIP, however, the Hfq-specific band was more intense and the signal-to-noise ratio was significantly higher. Based on these observations, one important benefit of PAR-CLIP is expected to be the reduced capture of non-specific targets. Additionally, as PAR-CLIP yielded higher levels of crosslinked RNA compared to standard CLIP, libraries made from RNA derived via PAR-CLIP are expected to be more comprehensive.

### Identification of Hfq-interacting RNAs using PAR-CLIP

To convert the crosslinked RNA into sequencing libraries, bands corresponding to the Hfq-crosslinked RNA were cut out from nitrocellulose strips and the RNA was released from the nitrocellulose strips after proteinase K treatment to digest protein. The purified RNAs were converted to sequencing libraries and sequenced on an Illumina HiSeq Sequencing system. To assess any contribution from background RNA, PAR-CLIP was similarly performed on CJ2109 with RNA purified from the same region of the nitrocellulose filters as was for FLAG-tagged Hfq. The resulting data were analyzed using the Galaxy bioinformatics platform ^13^ and the number of reads corresponding to each gene was determined.

To determine the utility of PAR-CLIP, several different criteria were used. First, the fraction of PAR-CLIP sequence reads from CJ2192 that correspond to the different RNA categories was determined. For normalization purposes, libraries made from total RNA preparations were also sequenced (RNA-Seq). Among the four major types of cellular RNAs (mRNAs, rRNAs, tRNAs and sRNAs), most of the uniquely mapping reads in the RNA-Seq samples were found to correspond to the first three categories, with the remainder (2.7%) being sRNAs (Fig. 3A). In the PAR-CLIP library, the reads for mRNAs, rRNAs and tRNAs varied moderately, but the fraction of sRNAs reads increased to 25%. The sharp increase in the sRNA reads is in line with expectations, since the major role of Hfq inside the cell is to bind to sRNAs and deliver them to their mRNA targets.

**Figure 3.**
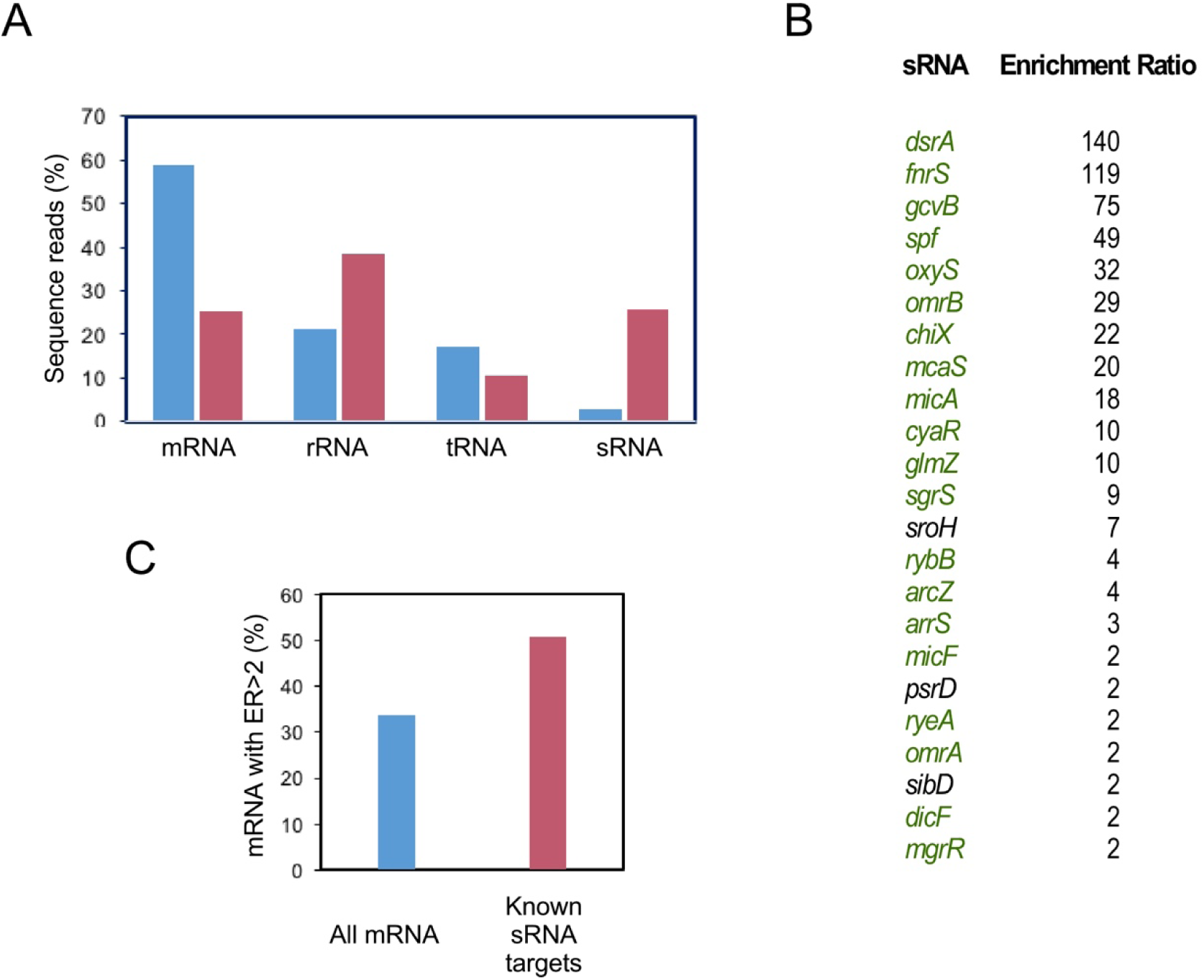
Analysis of Hfq PAR-CLIP data. (A) PAR-CLIP on Hfq enriches for sRNAs. The percentage of reads from CJ2192 that correspond to the different categories of *E. coli* RNAs are depicted by blue bars (Total RNA-Seq) or red bars (PAR-CLIP Seq). (B) sRNAs enriched in Hfq PAR-CLIP libraries. The sRNAs with ER values ≥ 2 are listed; those that correspond to previously identified Hfq-binding targets are colored green. (C) Comparison of PAR-CLIP-enriched mRNAs and validated Hfq-regulated mRNAs. The fraction of the mRNAs in PAR-CLIP libraries with ER ≥ 2 are depicted by a blue bar. The fraction of known mRNA targets of Hfq-binding sRNAs with ER ≥ 2 are indicated by a red bar.

Next, to evaluate the effectiveness of PAR-CLIP, we used a measure, termed “enrichment ratio” (ER) that corresponds to the relative fraction of reads for any RNA in the PAR-CLIP library prepared from CJ2192 relative to the corresponding library from CJ2109. A similar parameter has been used for Hfq in RIP-Seq experiments performed in *Salmonella typhimurium* ^14^. To improve accuracy, only the RNAs that yielded non-zero reads in each of duplicate PAR-CLIP libraries were considered. This procedure, when applied to sRNAs, yielded a list of 23 sRNAs that displayed a two-fold or greater enrichment score (Fig. 3B). Notably, 20 of the 25 previously described sRNA targets of Hfq were found to be included on this list ^15^. These results indicate that 80% of known Hfq targets are captured through PAR-CLIP.

We also found that 1686 mRNAs were reproducibly identified in duplicate PAR-CLIP libraries, and of these 570 (34%) had an ER value ≥ 2 (Supplementary Table 1). As >100 mRNAs have been validated as Hfq-dependent targets of sRNAs ^16^, we asked whether such RNAs are captured by PAR-CLIP more frequently than by chance. The potential reasons for crosslinking of these mRNAs could be either direct binding to Hfq or indirect binding via an association with Hfq-associated sRNAs. Of the 71 mRNAs that are both known to be regulated through Hfq-sRNA interactions and were present in the PAR-CLIP library list, 36 (51%) had an ER value ≥2, (P<0.01), significantly more than the numbers expected by chance (Fig. 3C; Supplementary Table 2). Based on the correlation between the mRNAs that are regulated by Hfq and the mRNAs enriched in Hfq PAR-CLIP libraries, we expect that a deeper analysis of the mRNAs identified in the PAR-CLIP libraries will result in the identification of many more Hfq-regulated mRNAs than are known at present.

## Discussion

PAR-CLIP was developed less than 10 years ago, but within a relatively short time, it has been implemented for use both in higher and lower eukaryotes ^17–20^. For reasons that are unclear, this technique has not been applied to prokaryotic organisms as yet. To determine whether PAR-CLIP can be used to characterize prokaryotic RNA-BPs, several experiments were performed. First, we found that the addition of 4SU to the growth medium allows this nucleoside to be incorporated into RNA. When cells were grown with 300 μM of 4SU in the medium, the fraction of uridine encoded residues with 4SU was found to be 1.3%, close to the 1.4-2.4% range of 4SU incorporation that has been found for eukaryotic cells ^8^. Thus, the ability to apply PAR-CLIP in bacteria is not limited by the import of 4SU into cells or by discrimination against this base by prokaryotic RNA polymerase.

Next, to determine conditions suitable for implementing PAR-CLIP, we evaluated crosslinking efficiency in cells grown with different levels of 4SU. We found that the extent of crosslinking initially rose with increased 4SU concentration, but high 4SU concentrations caused a signal reduction. Concurrently, high 4SU concentrations were found to reduce the growth rate. The highest amount of crosslinking was observed with 500 μM 4SU in the medium, which therefore represents a concentration suitable for maximal crosslinking efficiency. Alternatively, if minimizing the effects of 4SU on cell growth is desired, PAR-CLIP can also be performed on cells grown with 250 μM 4SU effectively. Although we have not tested 4SU incorporation in other prokaryotic organisms, we expect that a similar approach will be useful to determine the optimum conditions for the use of PAR-CLIP in other organisms.

We applied PAR-CLIP to Hfq, and generated sequence libraries from RNA crosslinked to this protein, as well from total RNA. A comparison between total RNA and PAR-CLIP reads showed that the sRNA category exhibited the largest increase, affirming the central role of Hfq in sRNA function (Fig 3A). Furthermore, of the 23 sRNAs enriched in the Hfq libraries, 20 corresponded to known Hfq targets (Fig. 3B), and only five previously described Hfq targets (*ryhB*, *rydC*, *sdsR*, *gadY* and *rprA*) were not identified in this set. To the best of our knowledge, comparable data using standard UV-CLIP have not been described, but the 80% success rate for PAR-CLIP suggests that this method should be useful to identify the majority of the RNA targets for other RNA-BPs. Moreover, only three of the sRNAs present in the enriched fraction are not established Hfq targets, indicating a low level of false-positives regardless of whether these sRNAs are found to be Hfq targets in the future or not. A combination of a high true positive and low false positive rate, as observed with PAR-CLIP, provides confidence that the application of this method to other proteins will yield a substantially reliable list of genuine RNA targets.

An additional benefit of PAR-CLIP is the increased level of T>C conversion that is observed at RNA crosslinking sites. This fortuitous change is caused by misreading of crosslinked 4SU residues during the reverse transcription process and has been exploited to define interaction sites between RNA-BPs and their RNA targets. We have also observed widespread T>C changes in the sequencing data from Hfq PAR-CLIP libraries. Although we have not analyzed these data further, the possibility to do so remains an option for any RNA-BP analyzed by PAR-CLIP, which may be considered as an added benefit of this method.

## Methods

### Strains and plasmids

The wild-type strain used for this study is CJ2109, a derivative of MG1655 that contains a point mutation in *rph*, which restores the reading frame of this prematurely terminated gene ^21^. Plasmid pCJ1099 was constructed by amplifying chromosomal DNA using primers with the sequence 5’ AAAAAGCTTGTCATGGCAGGATTATTCATCG 3’ and 5’ TTGGATCCTACTTGTCATCGTCATCCTTGTAGTCGATGTCATGATCTTTATAATCACCGTCATGGTCTTTGTAGTCCTGCGCAGCGGCAGGTTTACGCGG 3’. The PCR product, which contains the RhlE gene and an appended 3X-FLAG-tag sequence, was digested with BamHI and HindIII and subcloned into the BamHI and HindIII-digested arabinose-inducible expression vector, pMPM-A4 ^22^. CJ2191 was made by transforming pCJ1099 into CJ2109. Strain CJ2192, was constructed by appending a 3X FLAG-tag sequence to the C-terminus of the chromosomal Hfq gene by using recombineering ^23^.

### Analysis of 4SU incorporation into RNA

CJ2109 was grown in LB medium in the presence or absence of 300 μM 4SU and total RNA was isolated from mid-log cells using the hot-phenol method. 15 μg of each RNA sample was digested to nucleosides using snake venom phosphodiesterase and alkaline phosphatase, and analyzed by HPLC, as described ^8^.

### PAR-CLIP

25 ml of bacterial culture was grown to an OD_600_ of 0.5 in the absence or presence of 4SU added at early log-phase. For experiments using CJ2191, arabinose was added to a final concentration of 0.2% at OD_600_ = 0.2. Cells were cooled rapidly by adding ice and pelleted in a centrifuge. The cell pellets were resuspended in 10 ml of ice cold saline solution and crosslinked at 365 nm on ice for five min, or the indicated times, using a CL-1000 ultraviolet crosslinker (UVP). The cells were re-pelleted by centrifugation and stored at −80°C. Thawed cells were resuspended in 200 μl NP‐T buffer (50 mM Sodium phosphate, 300 mM NaCl, 0.05% Tween, pH 8.0), sonicated and centrifuged in a microfuge at 4°C. The supernatant fraction was transferred to a new tube and an equal volume of NP-T buffer supplemented with 8 M urea was added. The supernatant was incubated at 65°C for five min, followed by the addition of additional NP-T buffer to yield a final urea concentration of 1 M. The lysate was added to pre-washed anti‐FLAG magnetic beads (15 μl of a 50% bead suspension; Sigma, cat # A2220) and the mixture was mixed with rotation for 1 h at 4°C. The beads were collected by centrifugation at 8000 × *g* for 30 sec, resuspended in 100 μl NP‐T buffer, and washed twice with 100 μl high‐salt buffer (50 mM NaH_2_PO_4_, 1 M NaCl, 0.05% Tween, pH 8.0) and twice with 100 μl NP‐T buffer. The beads were resuspended in 100 μl of NP‐T buffer containing 1 mM MgCl_2_ and 50 units of Benzonase (Sigma; cat # E1014) and incubated for 10 min at 37°C in a heat block. The beads were washed once with high‐salt NP‐T buffer, twice with CIP buffer (100 mM NaCl, 50 mM Tris–HCl pH 7.4, 10 mM MgCl_2_), resuspended in 100 μl CIP buffer with 10 units of calf intestinal alkaline phosphatase (NEB) and incubated for 30 min at 37°C in a heat block. The beads were washed once with high‐salt buffer, twice with PNK buffer (50 mM Tris–HCl, pH 7.4, 10 mM MgCl_2_, 0.1 mM spermidine), resuspended in 100 μl PNK buffer with 10 units of T4 polynucleotide kinase and 1 μCi of *γ*-^32^P-ATP, and incubated for 30 min at 37°C. The beads were washed thrice with NP-T buffer and resuspended in 20 μl Protein Loading buffer (62 mM Tris, pH 6.8, 2% SDS, 0.01% bromophenol blue, 10% glycerol) and incubated for 3 min at 95°C. The beads were pelleted by centrifugation, and the supernatant was loaded on a mini-Protean 4-15% SDS–polyacrylamide gel (Bio-Rad), along with pre-stained protein markers. The RNA–protein complexes were transferred to nitrocellulose membrane using a Mini Trans-blot electrophoretic transfer cell (BioRad). The protein marker bands were overlaid with radioactive ink, and the membrane was exposed overnight on a phosphorimager screen. The bands corresponding to labeled RNA–protein complexes were cut into small pieces and incubated at 37°C with 400 μl PK solution [50 mM Tris– HCl, pH 7.4, 75 mM NaCl, 6 mM EDTA, 1% SDS, 10 units of RNasin (Promega) and 400 μg of proteinase K (ThermoScientific)] for 30 min with shaking at 37°C. 100 μl of 9 M urea solution was added and the incubation was continued for an additional 30 min. 450 μl of the PK/urea solution was mixed with 450 μl phenol/chloroform/isoamyl alcohol (25:24:1) alcohol and centrifuged for 12 min (16,000 × *g*) at 4°C. The aqueous phase was precipitated by overnight incubation at −20°C after the addition of 3 volumes of ethanol, 1/10 volume of 3 M Sodium Acetate (pH 5.2) and 1 μl of GlycoBlue (Life Technologies). The precipitate was pelleted by centrifugation (30 min, 16,000 × *g*, 4°C), washed with 80% ethanol, centrifuged again (15 min, 16,000 × *g*, 4°C), dried and resuspended in 10 μl of sterile water.

### Sequence library preparation

High throughput sequencing libraries were prepared using RNA isolated from PAR-CLIP samples using the NEBNext Multiplex Small RNA Library Prep Set for Illumina (Set 2; cat# E7580, New England Biolabs) according to the manufacturer’s instructions. To prepare RNA-Seq libraries, 1 μg of total RNA was digested with 0.01 units of Benzonase for 10 min at 37°C in 20 μl of NP-T buffer, followed by extraction with phenol/chloroform/isoamyl alcohol and ethanol precipitation. 100 ng of the digested RNA was used for sequence library preparation as described above.

### Bioinformatics Data Analysis

Raw sequencing data were analyzed by using the web-based Galaxy platform ^13^. Raw data was converted to Fastqsanger format and adaptors were removed using Trim Galore! The reads were aligned to the *E. coli* genome (Genbank: NC_000913.3) using Bowtie. Uniquely aligned reads corresponding to individual RNAs were determined using the htseq-count tool. Reads from the mRNA 5’ UTR regions were separately counted using annotations based on the mapping of RNA 5’-ends ^24^ and combined with the reads for the corresponding open-reading frames. ER values were calculated as the geometric mean of the relative frequency of reads in duplicate PAR-CLIP libraries derived from the Hfq-tagged strain divided by the reads in duplicate PAR-CLIP libraries derived from the wild-type strain.

## Acknowledgements

We thank Dr. Zhongwei Li (Florida Atlantic University) for assistance with HPLC analysis. We thank Thorsten Bischler and Sung-Huan Yu (University of Würzburg) for providing a GFF file with 5’ UTR annotations. This work was supported by Grant GM114540 from the National Institutes of Health.

## Author Contributions

C. J. and S. O. designed the studies, performed the experiments and approved the manuscript.

