## Supplementary material for "Development of PAR-CLIP to analyze RNA-protein interactions in prokaryotes": supplemantal table 1

Supplementary Data for “Development of PAR-CLIP to analyze RNA-protein interactions in prokaryotes” by Sandeep Ojha and Chaitanya Jain

Supplementary Table 1: mRNAs enriched in Hfq PAR-CLIP libraries by ≥ 2-fold are listed. ER values for mRNAs that yielded non-zero PAR-CLIP reads from the Hfq-tagged strain but none from the wild-type strain are denoted by a hashtag.

| **Gene** | **ER** | **Gene** | **ER** | **Gene** | **ER** | **Gene** | **ER** | **Gene** | **ER** | **Gene** | **ER** |
| --- | --- | --- | --- | --- | --- | --- | --- | --- | --- | --- | --- |
| *guaD* | # | *araG* | # | *pyrE* | 14 | *ibaG* | 8.1 | *recD* | 6.3 | *yhaO* | 5.3 |
| *ygdR* | # | *yeaO* | # | *gatA* | 14 | *atpI* | 8.1 | *malY* | 6.3 | *putA* | 5.2 |
| *rhtC* | # | *tnaB* | 117 | *waaQ* | 14 | *yqiB* | 8.1 | *tdcG* | 6.3 | *nadR* | 5.2 |
| *nudG* | # | *fxsA* | 73 | *proY* | 13 | *hsrA* | 8.0 | *entS* | 6.2 | *feaB* | 5.2 |
| *ybiO* | # | *shiA* | 60 | *dsbB* | 13 | *mppA* | 7.9 | *ppx* | 6.2 | *gnsA* | 5.2 |
| *yfcG* | # | *mutS* | 48 | *trg* | 13 | *mprA* | 7.9 | *fliR* | 6.2 | *yedI* | 5.2 |
| *ltaE* | # | *waaS* | 42 | *araJ* | 13 | *yqhD* | 7.8 | *treA* | 6.2 | *cysE* | 5.2 |
| *cybB* | # | *trxC* | 35 | *ycjF* | 12 | *leuD* | 7.8 | *aroP* | 6.1 | *kup* | 5.2 |
| *flc* | # | *msrC* | 35 | *ghrA* | 12 | *yadH* | 7.7 | *rcsD* | 6.1 | *mrdB* | 5.2 |
| *ydcD* | # | *gtrA* | 35 | *uidR* | 12 | *gcvH* | 7.7 | *eutC* | 6.1 | *pntB* | 5.1 |
| *pgpB* | # | *waaZ* | 34 | *ibpA* | 12 | *ndk* | 7.6 | *mglB* | 6.0 | *ydhR* | 5.1 |
| *cdh* | # | *nrdF* | 31 | *sucD* | 12 | *gstB* | 7.6 | *yebO* | 6.0 | *yiaD* | 5.1 |
| *yebW* | # | *mqsA* | 31 | *metN* | 12 | *mnmE* | 7.6 | *fiu* | 6.0 | *wbbJ* | 5.1 |
| *yqeC* | # | *yfbU* | 29 | *yeaD* | 12 | *lnt* | 7.6 | *yigL* | 6.0 | *yghA* | 5.1 |
| *metC* | # | *tusB* | 29 | *lldP* | 12 | *pheL* | 7.5 | *ubiI* | 6.0 | *ydhQ* | 5.1 |
| *soxR* | # | *chiP* | 28 | *ilvC* | 11 | *yjjJ* | 7.5 | *cpdA* | 6.0 | *yhaJ* | 5.0 |
| *yciC* | # | *ribF* | 28 | *emrD* | 11 | *yjeI* | 7.3 | *sdhD* | 6.0 | *gatD* | 5.0 |
| *yfhH* | # | *marC* | 27 | *yhhX* | 11 | *fbp* | 7.3 | *wbbK* | 5.9 | *rbsD* | 5.0 |
| *yeiL* | # | *waaO* | 23 | *ygiQ* | 11 | *rsmC* | 7.3 | *dcyD* | 5.9 | *aqpZ* | 5.0 |
| *argR* | # | *gltI* | 22 | *lldD* | 11 | *pldB* | 7.2 | *folC* | 5.9 | *menF* | 4.9 |
| *yidQ* | # | *wzyE* | 22 | *rlmD* | 11 | *ycaO* | 7.2 | *mdfA* | 5.9 | *yiaF* | 4.9 |
| *yoaJ* | # | *yjhU* | 22 | *ydgK* | 11 | *exuT* | 7.1 | *yhbJ* | 5.9 | *rpsJ* | 4.9 |
| *rhsD* | # | *wzxB* | 21 | *rpoE* | 10 | *rbbA* | 7.1 | *putP* | 5.8 | *yajR* | 4.9 |
| *ampE* | # | *dhaK* | 21 | *serA* | 10 | *metB* | 7.0 | *ppiA* | 5.8 | *cmoB* | 4.9 |
| *insX* | # | *fepB* | 20 | *fruK* | 10 | *yoaE* | 6.9 | *yceM* | 5.8 | *yhfA* | 4.9 |
| *sgcB* | # | *pdxJ* | 20 | *hsdS* | 10 | *ubiA* | 6.9 | *yecR* | 5.8 | *wbbH* | 4.9 |
| *opgC* | # | *ispB* | 19 | *thrC* | 9.8 | *nadK* | 6.9 | *menA* | 5.8 | *glnA* | 4.9 |
| *htrE* | # | *pgpC* | 19 | *sufS* | 9.7 | *ygdQ* | 6.9 | *nanA* | 5.8 | *ppsA* | 4.9 |
| *icdC* | # | *yfjW* | 19 | *mdtK* | 9.6 | *pssA* | 6.9 | *aceK* | 5.8 | *glnE* | 4.9 |
| *ilvE* | # | *wbbL* | 18 | *glpG* | 9.5 | *plsB* | 6.8 | *sgrT* | 5.8 | *yedD* | 4.8 |
| *uspB* | # | *mglC* | 18 | *ddlB* | 9.3 | *dut* | 6.8 | *queC* | 5.8 | *ligA* | 4.8 |
| *panM* | # | *yidB* | 18 | *rseC* | 9.2 | *yohJ* | 6.8 | *srlE* | 5.7 | *mipA* | 4.7 |
| *yzgL* | # | *uhpT* | 18 | *wzxE* | 9.1 | *kdsA* | 6.7 | *uraA* | 5.7 | *rsmH* | 4.7 |
| *mcbA* | # | *narP* | 18 | *fre* | 9.0 | *rluB* | 6.7 | *mlaD* | 5.7 | *glpB* | 4.7 |
| *yhaI* | # | *ibpB* | 18 | *metL* | 9.0 | *cof* | 6.7 | *tonB* | 5.7 | *hsdM* | 4.6 |
| *gatR* | # | *yncL* | 18 | *mglA* | 9.0 | *ytfP* | 6.7 | *borD* | 5.6 | *yigA* | 4.6 |
| *ynbD* | # | *ydhC* | 17 | *yrfG* | 8.9 | *yjiT* | 6.6 | *ynfH* | 5.6 | *dacA* | 4.6 |
| *tag* | # | *yfhG* | 17 | *acrD* | 8.9 | *gtrS* | 6.6 | *mazG* | 5.6 | *gntR* | 4.6 |
| *yciF* | # | *bisC* | 17 | *carB* | 8.8 | *yfeS* | 6.6 | *ycbZ* | 5.6 | *mpl* | 4.5 |
| *ghxQ* | # | *yliF* | 17 | *prlC* | 8.8 | *cpxA* | 6.5 | *recF* | 5.6 | *thrL* | 4.5 |
| *ychQ* | # | *yjiV* | 16 | *yoaF* | 8.8 | *hsdR* | 6.5 | *sohB* | 5.5 | *tadA* | 4.5 |
| *xylF* | # | *yigM* | 15 | *sapB* | 8.7 | *tatC* | 6.5 | *potA* | 5.5 | *amn* | 4.5 |
| *hisH* | # | *yiiR* | 15 | *nmpC* | 8.5 | *dhaL* | 6.5 | *fliY* | 5.5 | *rpe* | 4.5 |
| *yqiG* | # | *alx* | 15 | *sapC* | 8.5 | *lrhA* | 6.4 | *tyrR* | 5.5 | *gatB* | 4.5 |
| *yjhB* | # | *uvrY* | 15 | *bcsE* | 8.5 | *hemG* | 6.4 | *tdcC* | 5.4 | *ydbJ* | 4.5 |
| *nanK* | # | *paaX* | 15 | *ompR* | 8.4 | *wecF* | 6.3 | *ygiM* | 5.4 | *ompF* | 4.5 |
| *yejE* | # | *gntX* | 15 | *miaB* | 8.2 | *ybhS* | 6.3 | *yeaP* | 5.4 | *exbB* | 4.5 |
| *thiK* | # | *lldR* | 15 | *yihY* | 8.1 | *wbbI* | 6.3 | *ybgJ* | 5.3 | *sstT* | 4.5 |
| *pspB* | # | *metI* | 14 | *waaR* | 8.1 | *yafV* | 6.3 | *yoaA* | 5.3 | *mcrB* | 4.5 |
| *paaJ* | # | *alaC* | 14 | *ypjD* | 8.1 | *yqeF* | 6.3 | *artQ* | 5.3 | *rsmA* | 4.4 |

| **Gene** | **ER** | **Gene** | **ER** | **Gene** | **ER** | **Gene** | **ER** | **Gene** | **ER** | **Gene** | **ER** |
| --- | --- | --- | --- | --- | --- | --- | --- | --- | --- | --- | --- |
| *yajC* | 4.4 | *gabT* | 3.5 | *yidE* | 3.0 | *ybaY* | 2.6 | *uvrD* | 2.3 | *yjtD* | 2.1 |
| *lrp* | 4.4 | *sucC* | 3.5 | *mtn* | 3.0 | *ychA* | 2.6 | *yfbT* | 2.3 | *icd* | 2.1 |
| *yebZ* | 4.4 | *yciM* | 3.5 | *nudE* | 2.9 | *cyoD* | 2.6 | *suhB* | 2.3 | *argF* | 2.1 |
| *yiaG* | 4.4 | *cysM* | 3.5 | *secB* | 2.9 | *lspA* | 2.6 | *metG* | 2.3 | *hns* | 2.1 |
| *glpX* | 4.3 | *motB* | 3.5 | *thiB* | 2.9 | *ftsE* | 2.6 | *gltB* | 2.3 | *slyD* | 2.1 |
| *yecS* | 4.3 | *ychN* | 3.5 | *ompX* | 2.9 | *rfbC* | 2.6 | *mscS* | 2.3 | *hfq* | 2.1 |
| *nusB* | 4.3 | *sdiA* | 3.5 | *asnB* | 2.9 | *tolC* | 2.6 | *alaS* | 2.3 | *secD* | 2.1 |
| *yfgM* | 4.3 | *cra* | 3.5 | *murI* | 2.9 | *serC* | 2.6 | *secF* | 2.3 | *ymdA* | 2.1 |
| *yqfA* | 4.3 | *pqiA* | 3.5 | *kefB* | 2.9 | *dedA* | 2.6 | *flgN* | 2.3 | *hslV* | 2.0 |
| *cirA* | 4.2 | *rnd* | 3.4 | *ypfN* | 2.9 | *yraQ* | 2.6 | *wecB* | 2.3 | *prmB* | 2.0 |
| *alr* | 4.2 | *rlmM* | 3.4 | *malT* | 2.9 | *dtpD* | 2.6 | *ydhF* | 2.3 | *dnaE* | 2.0 |
| *fadJ* | 4.2 | *dppD* | 3.4 | *sdhC* | 2.9 | *maeA* | 2.6 | *nlpI* | 2.3 | *tsr* | 2.0 |
| *dnaA* | 4.2 | *yfcA* | 3.4 | *srlD* | 2.9 | *nanT* | 2.6 | *waaB* | 2.3 | *dctA* | 2.0 |
| *yhjG* | 4.1 | *proS* | 3.4 | *tnaC* | 2.9 | *ptsG* | 2.6 | *gpmM* | 2.3 | *degP* | 2.0 |
| *nfsB* | 4.1 | *tpx* | 3.3 | *plsC* | 2.9 | *nuoN* | 2.6 | *srkA* | 2.3 | *mlaB* | 2.0 |
| *prpC* | 4.1 | *yhbS* | 3.3 | *metH* | 2.9 | *nlpC* | 2.5 | *yhcN* | 2.3 | *tatA* | 2.0 |
| *glmS* | 4.1 | *cycA* | 3.3 | *bioH* | 2.9 | *nhaA* | 2.5 | *mepM* | 2.3 | *rnlA* | 2.0 |
| *glpF* | 4.1 | *carA* | 3.3 | *argG* | 2.9 | *yjiY* | 2.5 | *aceB* | 2.2 | *trxB* | 2.0 |
| *lpxT* | 4.0 | *dtd* | 3.3 | *yjiR* | 2.9 | *envZ* | 2.5 | *yohP* | 2.2 | *tolR* | 2.0 |
| *modF* | 4.0 | *pnuC* | 3.3 | *alaE* | 2.8 | *fepD* | 2.5 | *cysU* | 2.2 | *yobF* | 2.0 |
| *ypfM* | 4.0 | *kgtP* | 3.3 | *mgtA* | 2.8 | *zipA* | 2.5 | *bamD* | 2.2 |  |  |
| *sthA* | 4.0 | *rpmE* | 3.3 | *yfbS* | 2.8 | *ymdB* | 2.5 | *ylaC* | 2.2 |  |  |
| *dsbD* | 4.0 | *nlpD* | 3.3 | *yfeX* | 2.8 | *mukF* | 2.5 | *miaA* | 2.2 |  |  |
| *rbsB* | 4.0 | *tnaA* | 3.3 | *osmC* | 2.8 | *yhbW* | 2.5 | *glgB* | 2.2 |  |  |
| *dmlR* | 4.0 | *clpP* | 3.3 | *clpS* | 2.8 | *yshB* | 2.5 | *yedE* | 2.2 |  |  |
| *yahN* | 3.9 | *greB* | 3.3 | *fabZ* | 2.8 | *gntU* | 2.5 | *nuoH* | 2.2 |  |  |
| *gmhA* | 3.9 | *ydgD* | 3.3 | *maeB* | 2.8 | *tgt* | 2.5 | *rpoS* | 2.2 |  |  |
| *eptB* | 3.8 | *yaeH* | 3.3 | *yhfK* | 2.8 | *rnc* | 2.5 | *aroG* | 2.2 |  |  |
| *tatE* | 3.8 | *iscU* | 3.3 | *grxC* | 2.8 | *ychJ* | 2.5 | *ybjJ* | 2.2 |  |  |
| *wecC* | 3.8 | *yghB* | 3.2 | *flgL* | 2.7 | *zapE* | 2.5 | *ybjX* | 2.2 |  |  |
| *treC* | 3.8 | *mtfA* | 3.2 | *sppA* | 2.7 | *sufC* | 2.5 | *sucA* | 2.2 |  |  |
| *flgA* | 3.8 | *flgE* | 3.2 | *barA* | 2.7 | *moeA* | 2.5 | *gcvA* | 2.2 |  |  |
| *dcuS* | 3.8 | *amiA* | 3.2 | *ampG* | 2.7 | *recG* | 2.5 | *gtrB* | 2.2 |  |  |
| *cysS* | 3.7 | *murJ* | 3.2 | *mraZ* | 2.7 | *glnG* | 2.4 | *yncD* | 2.2 |  |  |
| *murF* | 3.7 | *sseA* | 3.2 | *racR* | 2.7 | *rodZ* | 2.4 | *flgB* | 2.2 |  |  |
| *phoR* | 3.7 | *rnpA* | 3.1 | *hslU* | 2.7 | *pntA* | 2.4 | *yhiI* | 2.2 |  |  |
| *yqjA* | 3.7 | *nhaR* | 3.1 | *mlaF* | 2.7 | *mtlR* | 2.4 | *yaiE* | 2.2 |  |  |
| *ftsQ* | 3.7 | *hupB* | 3.1 | *glpK* | 2.7 | *waaG* | 2.4 | *fhuA* | 2.1 |  |  |
| *yffS* | 3.6 | *glpT* | 3.1 | *yfeH* | 2.7 | *osmF* | 2.4 | *ldtD* | 2.1 |  |  |
| *yjiA* | 3.6 | *rbsC* | 3.1 | *rsxA* | 2.7 | *fliC* | 2.4 | *cpxP* | 2.1 |  |  |
| *brnQ* | 3.6 | *napA* | 3.1 | *plsY* | 2.7 | *mltB* | 2.4 | *sucB* | 2.1 |  |  |
| *nuoA* | 3.6 | *opgG* | 3.0 | *ptsP* | 2.7 | *tyrS* | 2.4 | *rna* | 2.1 |  |  |
| *gdhA* | 3.6 | *proQ* | 3.0 | *glmU* | 2.7 | *yeaQ* | 2.4 | *gapC* | 2.1 |  |  |
| *pepP* | 3.6 | *gatC* | 3.0 | *nuoB* | 2.7 | *minC* | 2.4 | *metK* | 2.1 |  |  |
| *nupC* | 3.6 | *ybjL* | 3.0 | *ftsI* | 2.7 | *tolB* | 2.4 | *dxs* | 2.1 |  |  |
| *aroA* | 3.6 | *mhpR* | 3.0 | *trkG* | 2.7 | *ftsW* | 2.4 | *ptrB* | 2.1 |  |  |
| *dgt* | 3.6 | *fliZ* | 3.0 | *bepA* | 2.7 | *flhB* | 2.4 | *yicC* | 2.1 |  |  |
| *flgD* | 3.6 | *tsx* | 3.0 | *tamA* | 2.7 | *mfd* | 2.3 | *yjiN* | 2.1 |  |  |
| *ygdH* | 3.6 | *oppA* | 3.0 | *glgC* | 2.7 | *lepB* | 2.3 | *yehU* | 2.1 |  |  |
| *mepS* | 3.6 | *cyoC* | 3.0 | *nupG* | 2.6 | *mreB* | 2.3 | *tsaB* | 2.1 |  |  |

Supplementary Table 2: ER values for validated targets of sRNAs.

| **Gene** | **ER** |
| --- | --- |
| *xylF* | # |
| *shiA* | 60.08 |
| *mutS* | 48.46 |
| *chiP* | 27.95 |
| *ompR* | 8.37 |
| *ygdQ* | 6.88 |
| *fiu* | 6.04 |
| *yigL* | 6.03 |
| *sdhD* | 5.96 |
| *ygiM* | 5.42 |
| *cysE* | 5.16 |
| *ompF* | 4.47 |
| *sstT* | 4.46 |
| *lrp* | 4.42 |
| *cirA* | 4.22 |
| *glmS* | 4.08 |
| *glpF* | 4.07 |
| *sthA* | 4.01 |
| *rbsB* | 3.98 |
| *eptB* | 3.83 |
| *gdhA* | 3.61 |
| *sucC* | 3.53 |
| *tpx* | 3.35 |
| *cycA* | 3.35 |
| *tsx* | 3.00 |
| *oppA* | 2.98 |
| *ompX* | 2.93 |
| *sdhC* | 2.87 |
| *mraZ* | 2.71 |
| *maeA* | 2.57 |
| *nanT* | 2.56 |
| *ptsG* | 2.56 |
| *rpoS* | 2.19 |
| *icd* | 2.09 |
| *hns* | 2.06 |
| *yobF* | 2.00 |
| *fur* | 1.99 |
| *sdhA* | 1.96 |
| *lamB* | 1.93 |
| *fadL* | 1.83 |
| *caiA* | 1.79 |
| *erpA* | 1.78 |
| *dppA* | 1.69 |
| *ydaM* | 1.67 |
| *ftsZ* | 1.61 |
| *ompC* | 1.59 |
| *sodA* | 1.54 |
| *ompT* | 1.52 |
| *ompT* | 1.52 |
| *fumA* | 1.40 |
| *yifK* | 1.30 |
| *iscR* | 1.20 |
| *cydD* | 1.12 |
| *nirB* | 1.08 |
| *sodB* | 1.04 |
| *manZ* | 1.01 |
| *sdaC* | 1.00 |
| *tig* | 0.95 |
| *gltA* | 0.86 |
| *lpp* | 0.86 |
| *acnA* | 0.83 |
| *hinT* | 0.79 |
| *cfa* | 0.78 |
| *iscS* | 0.69 |
| *ptsI* | 0.59 |
| *ptsI* | 0.59 |
| *gpmA* | 0.57 |
| *ompA* | 0.57 |
| *manY* | 0.32 |
| *msrB* | 0.24 |
| *fhlA* | 0.02 |
